# Evidence of long-term purging of mutation load in killer whale genomes

**DOI:** 10.1101/2021.08.21.457186

**Authors:** Andrew D. Foote, M. Thomas P. Gilbert, Shyam Gopalakrishnan, Marie Louis, Michael D. Martin, Phillip A. Morin, Milaja Nykänen, Camilla Hjorth Scharff-Olsen, Laurent Excoffier

## Abstract

Genomes of high latitude killer whales harbour signatures of post-glacial founding and expansion. Here, we investigate whether reduced efficacy of selection increased mutation load in founder populations, or whether recessive deleterious mutations exposed to selection in homozygous genotypes were purged. Comparing the accumulation of synonymous and non-synonymous mutations across pairs of globally sampled genomes reveals that the most significant outliers are high latitude North Atlantic genomes, which have accumulated significantly fewer non-synonymous mutations than all other populations. Comparisons with the genome of a 7.5-Kyr-old North Atlantic killer whale, inferred to be closely related to the population directly ancestral to present-day Icelandic and Norwegian populations, calibrates the timing of the action of selection on non-synonymous mutations predominantly to the mid-late Holocene. Non-synonymous mutations purged in modern Norwegian killer whale genomes are found as globally shared standing variation in heterozygote genotypes more often than expected, suggesting overdominance. Taken together, our findings are consistent with purging of recessive non-synonymous mutations exposed to selection in founder-associated homozygous genotypes.

---

Genetic variation among populations is influenced by the complex interaction between mutation, selection, gene flow and drift (Hahn 2019), which are in-turn influenced by demographic history (Charlesworth 2009). Mutations are a source of biological variation drawn from a distribution of fitness effects (Eyre-Walker and Keightley 2007). The term mutation load refers to the deleterious variants within an individual or population (Kimura et al. 1963; Henn et al. 2015). The frequency distribution of deleterious mutations within a population will vary among populations due to the respective effects of genetic drift and migration and the efficacy of purifying selection associated with demographic history. For example, weakly deleterious mutations can evolve nearly neutrally in small populations, and at the wave front of range expansions (Ohta 1973; Peischl et al. 2013; Lohmueller 2014; Sousa et al. 2014; Peischl and Excoffier 2016), consequently rising to high frequency (so-called gene surfing; Hallatschek and Nelson 2008; Excoffier et al. 2009). Modelling the effect of range expansions on standing genetic variation has demonstrated that while the number of deleterious mutations may remain unchanged, the frequency of deleterious recessive mutations in homozygous genotypes may increase (Peischl and Excoffier 2016). Whilst this process can increase deleterious recessive mutation load, it can also increase rates of purging of these same mutations by exposing variants to purifying selection once they are in homozygous genotypes (Kirkpatrick and Jarne 2000; Glemin 2003; Balick et al. 2015; Hedrick and Garcia-Dorado 2016). Despite these theoretical expectations, there is limited empirical evidence for purging due to recent demographic bottlenecks in wild populations (but see Xue et al. 2015; Robinson et al. 2018; Grossen et al. 2020; Stoffel et al. 2021). Given that range shifts are a common response to ongoing global warming (Chen et al. 2011), understanding how range expansions and shifts influence the fitness of natural populations at the range margins presents an increasingly important consideration for conservation (Gibson et al. 2009). Therefore, retrospectively investigating changes in the genetic composition of populations associated with range shifts in response to past climate change may provide important predictions of genetic changes associated with the ongoing climatic change (Díez-del-Molino et al. 2018).

Here, we explore mutation load variation associated with different demographic histories through analysis of whole genome sequences from a global dataset of modern killer whales and an ancient (7.5 Kyr old) killer whale (Figure 1a). Killer whales are the most widely distributed marine mammal, being found from the Arctic to the Antarctic and across all latitudes in between. The scale of their dispersal range means that much standing genetic variation (SGV) is shared globally (Foote et al. 2019), and shared SGV has been hypothesised to be maintained through episodic gene flow between demes (Foote et al. 2021). Genetic differentiation is greatest at the high latitude range margins, in waters that were covered by sea ice during the Last Glacial Maximum, consistent with range expansion into these waters during the Holocene (Foote et al. 2019, 2021). Thus, whilst modern humans typically show a clinal change in genetic diversity associated with expansion out of Africa (Henn et al. 2016), killer whale genomes suggest independent post-glacial range expansions in different ocean basins (Morin et al. 2015; Foote et al. 2016, 2019, 2021). Therefore, a comparison among globally distributed killer whale genomes offers the possibility to explore the genetic outcomes of independent, parallel demographic histories. Killer whales thus represent a useful comparative system to modern humans for studying the long-term relationship between demographic history, range expansions and individual mutation load.

**Figure 1:**
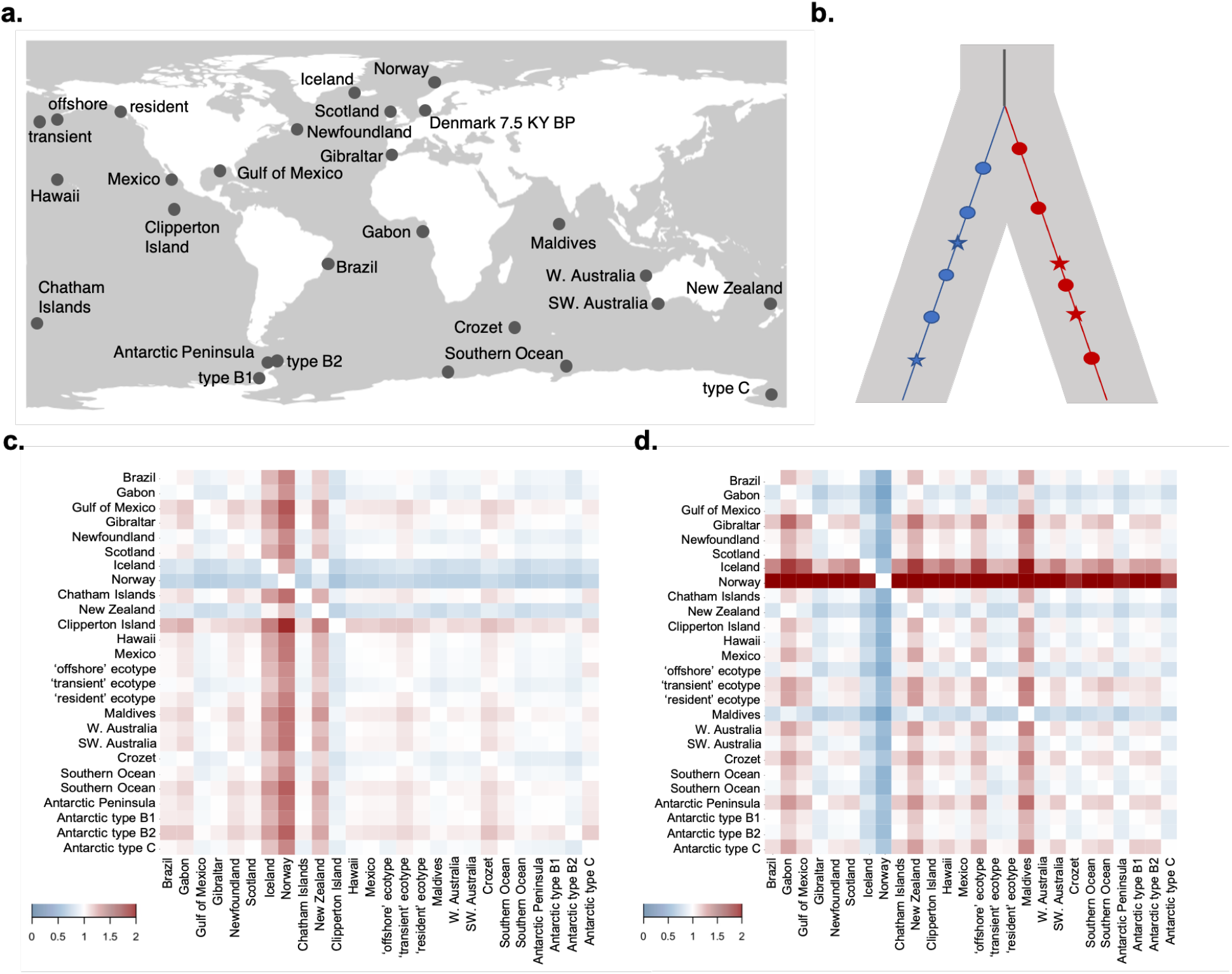
**a**. Sampling locations for the individuals included in this study. **b**. A schematic of the *R*_*X*/*Y*_ statistic which is derived from counts of mutations not shared by randomly sampled haploid genomes from two different populations *X* and *Y* (coloured blue and red), which have accumulated since they diverged from a shared common ancestor. Non-synonymous mutations are shown as stars, synonymous mutations as circles. Under neutral expectations, both classes of mutation should accumulate at an approximately equal rate in each population. (**c** & **d**) Heatmaps of *R*_*X*/*Y*_ estimates: the ratio of counts of mutations not shared by randomly sampled haploid genomes from population *X* (*x*-axis) and population *Y* (*y*-axis) accumulating in each genome since they diverged from a shared common ancestor, for **(c)** synonymous mutations, and **(d)** non-synonymous mutations.

## Results and Discussion

Killer whales are found in the highest density at high latitudes (Forney & Wade 2006), but effective population size (*N*_e_) is typically lower in high latitude populations, and highest at low latitudes (Foote et al. 2021). The efficacy of natural selection to remove deleterious mutations is dependent upon the product of the effective population size (*N*_e_) and the selection coefficient of the mutation (*s*) (Charlesworth 2009). Thus, a reduction in *N*_e_ among killer whale populations should be reflected in reduced efficacy of purifying selection (*N*_e_*s*) to remove deleterious mutations of equivalent selection coefficients (*s*) in different populations. However, the genomes of high latitude populations of killer whales typically harbour large numbers of short (<1Mb) runs of homozygosity (ROH), reflecting high background relatedness due to coalescence of haplotypes deep within the pedigree (Foote et al. 2021). Thus, many recessive deleterious mutations are expected to have been in homozygous genotypes and therefore exposed to selection for hundreds of generations.

To test for variation in the efficacy of selection among killer whale populations, we estimated the *R*′_*X*/*Y*_ statistic (Do et al. 2015; Figure 1b), which draws inference from counts of mutations not shared by randomly sampled haploid genomes from two different populations (*X* and *Y*). Under neutral expectations and an assumption of no inter-population variation in mutation rate, mutations should accumulate at an approximately equal rate in each population since they diverged from a shared common ancestor. However, differences in population size, and therefore *N*_e_*s*, between populations will result in an asymmetry in the accumulation, and removal by selection of deleterious mutations in genomes sampled from the differing populations. Genomes sampled from populations with smaller population size would thus be expected to harbour more deleterious mutations in their genomes, due to weaker purifying selection.

We first randomly selected an allele from each site of the genome of 26 killer whales sampled across their known range (Foote et al. 2019, 2021); generating a haploid genome for each population. We then measured the accumulation of derived non-synonymous relative to synonymous mutations per-genome across all possible pairs of populations from a global dataset. The ancestral state allele at each site was determined from the consensus of genomes of two species closely related to the killer whale (approximately 10 Myr since divergence, McGowen et al. 2020), the bottlenose dolphin *Tursiops truncatus* and long-finned pilot whale *Globicephala melas*. While recognising that the neutral evolution of all synonymous mutations is an unrealistic assumption (Kern and Hahn 2018; Ragsdale et al. 2018; Machado et al. 2018), non-synonymous mutations are mainly deleterious, reducing the fitness of carriers and their offspring (Eyre-Walker and Keightley 2007). We therefore expect, that non-synonymous mutations will typically evolve under stronger selective constraints than synonymous mutations. We furthermore only considered transversions to avoid including C→T and A→G misincorporations caused by post-mortem DNA damage through deamination of cytosine to uracil (Pääbo 1989) in the ancient genome. Note that non-synonymous transversions may generally be more deleterious than non-synonymous transitions (Zhang 2000; but see Zou & Zhang 2020).

We counted derived synonymous transversions found in genome *X* but not in genome *Y* (*L*_*X*/*Y*_), and vice versa (*L*_*Y*/*X*_); *R*_*X*/*Y*_ (synonymous) is then the ratio ((*L*_*X*/*Y*_)/(*L*_*Y*/*X*_)) of the two counts (Do et al. 2015). We then did the same for derived non-synonymous transversions, resulting in the statistic *R*_*X*/*Y*_ (non-synonymous). Under neutral expectations, *R*_*X*/*Y*_ is expected to equal 1; under differences in selection and therefore mutation load *R*_*X*/*Y*_ will be greater or less than 1, if *X* or *Y* respectively have a higher non-synonymous mutation load (Do et al. 2015). We found that the genomes sampled from Iceland and Norway had estimates of *R*_*X*/*Y*_ (synonymous) significantly higher than 1 in all pairwise comparisons (*Z*-score > 4; Figure 1c). In contrast, *R*_*X*/*Y*_ (non-synonymous) was significantly less than 1 in the same pairwise comparisons (*Z*-score > 4; Figure 1d). The Maldives genome had the highest accumulation of non-synonymous relative to synonymous alleles (*Z*-score > 4; Figure 1c,d).

The interpretation of the relative differences in counts of synonymous and non-synonymous derived mutations as reflecting differences in purifying selection assumes additivity and ignores dominance. Given that more deleterious mutations are typically also more recessive (Huber et al. 2018), and that model predictions are that changes in mutation load associated with range expansion will result from deleteriously recessive standing variation being exposed in homozygous genotypes (Peischl and Excoffier 2016), we sought to infer whether purged alleles were globally found in heterozygote genotypes more often than expected by chance, suggesting overdominance. We tested each site for heterozygote excess in the full dataset (acknowledging that our dataset is comprised of multiple populations and thus does not conform to the Hardy Weinberg assumptions, but note the Wahlund effect is expected to reduce heterozygosity relative to HWE expectations, which should thus lead to a conservative test); comparing derived non-synonymous transversions found in the Maldives genome but not the Norway genome (and thus candidates for having been purged by selection in the ancestors of the Norway genome), with derived non-synonymous transversions shared by the Maldives and Norway genomes. Only a single derived non-synonymous transversion shared between the Maldives and Norway genomes showed significant (P<0.001) heterozygote excess (inbreeding coefficient *F*_IS_ <. −0.5) in the global dataset, whilst nine derived non-synonymous transversions found in the Maldives genome but not shared with the Norway genome showed an excess of heterozygous genotypes (P<0.001 to P<0.0000001). Whilst heterozygote excess can also reflect genotyping errors at sites with poor mapability, the finding of these variants in multiple individuals, combined with our data filtering pipeline, makes it unlikely that this result reflects erroneous genotype calls.

Based on the observation that deleterious derived mutations are typically rare (*i*.*e*. at low frequency) within populations and are therefore typically in heterozygous genotypes in outbred populations (under Hardy-Weinburg expectations), we expect fitness effects to be determined by dominance and selection coefficients (1−*hs*). However, in inbred populations, deleterious derived mutations will be predominantly in homozygous genotypes with fitness (1−*s*), and thus less influenced by dominance (Huber et al. 2018). Taken together, the results are consistent with deleterious recessive alleles being neutrally harboured in heterozygote genotypes in low latitude populations such as the Maldives population. Recessive deleterious alleles are inferred to have been purged from the high latitude Iceland and Norway population after a combination of drift during the inferred ancestral bottleneck in this population (Foote et al. 2021) and then being exposed to selection in homozygous genotypes.

To better understand the timing of the apparent purging of non-synonymous mutations, we additionally compared modern genomes with the genome of an ancient 3×coverage killer whale genome sequenced from a subfossil found in Denmark. The ancient killer whale was carbon-dated to 7,530-7,780 yr BP (Aaris-Sørensen et al. 2010) and we estimate it to be closely related to the population directly ancestral to the present-day Iceland and Norway populations (Figure 2). We estimated the statistic *R’*_*X*/*Y*_ = *R*_*X*/*Y*_ (non-synonymous)/*R*_*X*/*Y*_ (synonymous) (Do et al. 2015). This statistic normalizes counts of non-synonymous mutations private to each genome by the counts of private synonymous mutations. The normalization corrects for the expectation that modern samples will have accumulated more mutations than the ancient sample due to less time having elapsed since the ancient individual diverged from a common ancestor with the modern individual (Do et al. 2015). Under neutral expectations, *R*′_*X*/*Y*_ is expected to equal 1; under differences in selection and therefore mutation load *R*′_*X*/*Y*_ will be greater or less than 1, if *X* or *Y* respectively have a higher non-synonymous mutation load (Do et al. 2015). For example, *R*′_*Norway*/*Y*_ and *R*′_*Iceland*/*Y*_ were significantly less than 1 (*Z*-score > 4) in all pairwise comparisons (Figure 3). *R*′_*Ancient*/*Y*_ was also less than 1 in all pairwise comparisons, except when *Y* was either the modern Iceland or Norway genome (Figure 3). These comparisons between the ancient and modern killer whale genomes suggest purging of non-synonymous mutations by purifying selection was already happening in the ancestral North Atlantic population as represented in this study by the ancient genome, during the early Holocene, and that it continued in subsequent generations, as shown by the smaller proportion of non-synonymous mutations in the modern Norway and Iceland genomes than the ancient genome, Our results suggest purging of non-synonymous mutation by purifying selection is thus a long-term process that continued throughout the Holocene.

**Figure 2:**
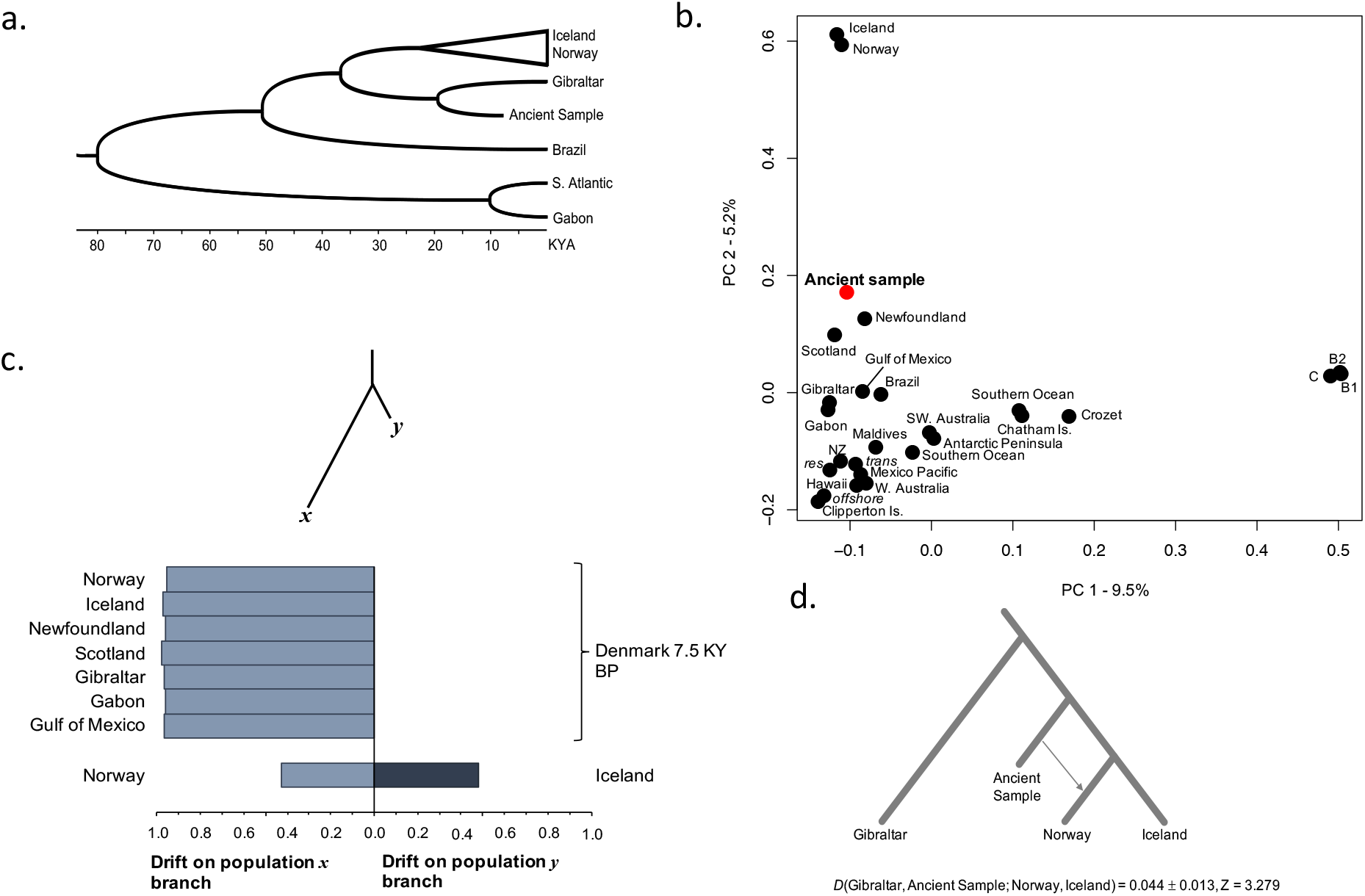
Relationship of the ancient (7,530-7,780 yr BP) killer whale to modern killer whale genomes. **a**. Cartoon subtree showing the mitochondrial haplotypes with the closest TMRCA to the ancient sample. Full Bayesian phylogenetic tree of 159 unique haplotype sequences generated from 453 samples is included in the supplementary materials. **b**. Principal Component Analysis (PCA) plot of the relationship of the ancient to the modern dataset based upon nuclear genomes. **c**. Site patterns and drift estimates for North Atlantic populations (*x*) versus the ancient sample (*y*), and at the bottom, for the modern Norway and modern Iceland samples to show how the test estimates approximately equal branch lengths from two modern samples. The estimate suggests the ancient sample pre-dates much of the drift that differentiates the modern Atlantic samples and is approximately equally ancestral to all. However, due to the low coverage, the estimated drift along branch *y* to the ancient sample will be downwardly biased. **d**. Testing of the hypothesis that the ancient sample is ancestral to the Norway and Iceland samples. D-statistic tests suggest the data are relatively consistent with the simple tree-like relationship represented above. The Norway and Iceland samples share a recent ancestor that diverged from an older shared ancestor with the ancient sample. However, there is a slight but significant (Z > 3) excess of shared alleles between the ancient and Norway samples, relative to between the ancient and Iceland samples. Thus, the relationship is slightly more complex than can be represented by a simple bifurcating model. The inclusion of the ancient sample in the ingroup with either the Norway or Iceland samples was rejected (Z > 30).

**Figure 3:**
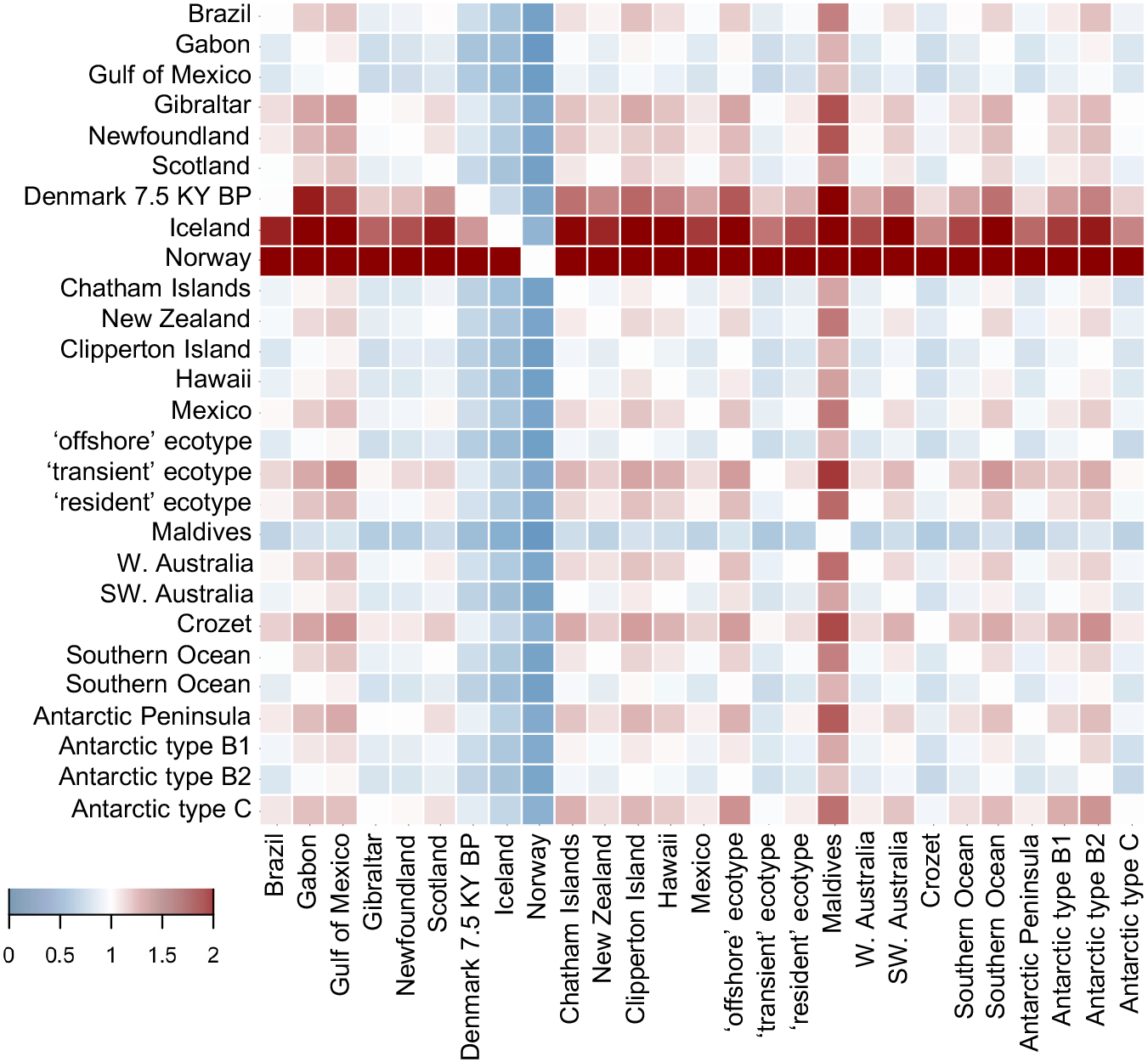
Relative load of non-synonymous mutations *R*′_*X*/*Y*_ for pairs of populations. Values greater than 1 (shaded red) indicate a higher proportion of non-synonymous to synonymous unshared derived mutations in population *X* (labelled on *x*-axis) than in population *Y* (labelled on *y*-axis); values less than 1 (shaded blue) indicate a higher proportion of synonymous to non-synonymous unshared derived mutations in population *X* than in population *Y*.

The Norwegian killer whale sample used in the *R*′_*X*/*Y*_ estimates was also the same individual used to generate the reference genome assembly (GCA_000331955.2; Foote et al. 2015). To determine whether reference bias (see Günther and Nettelblad 2019) had influenced our results, we generated a new reference fasta file from a pseudo-haploid North Pacific transient killer whale genome; which is highly divergent from the Icelandic and Norwegian genomes (Foote et al. 2019). The short-read data of the ancient killer whale, and four additional unrelated Norwegian killer whales sequenced for this study were then mapped to this new reference and *R*′_*Ancient*/*Y*_ was estimated as before. We found that *R*_*Norway*/*Y*_ (synonymous), but not *R*_*Norway*/*Y*_ (non-synonymous), were marginally upwardly biased when mapping to the Norwegian killer whale reference genome versus mapping to the Pacific transient reference genome. Nethertheless, for each of the modern Norwegian killer whales, *R*′_*Modern*/*Ancient*_ was also significantly less than 1 (*Z*-score > 4).

If the process underlying the apparent reduced non-synonymous mutation load in the high latitude North Atlantic population is the removal of deleterious recessive non-synonymous standing variation exposed to purifying selection in homozygous genotypes during range expansions, this begs the question, why do we not see a similar pattern in the peripheral populations of the North Pacific or Antarctic. Prior studies (Morin et al. 2015; Foote et al. 2021) have found that high latitude populations in these ocean basins show genetic characteristics of leading edge populations (Hewitt 2020), namely the formation of monophyletic clades and harbouring short runs of homozygosity (ROH) consistent with coalescence of identical by descent haplotypes deep within the pedigree and subsequent generations of recombination. However, there are differences between the North Pacific, Antarctic and North Atlantic. Admixture upon secondary contact between divergent lineages with different genetic backgrounds has been reported in both the North Pacific and Antarctic populations (Foote et al. 2019, 2021). Such admixture can break up ROH and ‘rescue’ inbred populations from recessive alleles, resulting in overdominance (Crow 1948; Bierne et al. 2002; Ingvarsson & Whitlock 2000; Whitlock et al. 2000). In contrast, all North Atlantic genomes sequenced to date coalesce in a shared common ancestor (Foote unpublished data), reducing the opportunity for heterosis.

What are the implications for the conservation of populations of killer whales and more broadly any species that will shift their range towards higher latitudes under current directional global warming? Our results largely agree with simulation results and the empirical findings (Kyriazis et al. 2021; Stoffel et al. 2021; van der Valk et al. 2021), that over the long-term, selection can remove deleterious mutations from small populations. However, the lack of a parallel pattern in other high latitude killer whale populations illustrates the complexity of these dynamics. Ultimately, predictive models of mutation load require a joint distribution of selection and dominance coefficients and a realistic demographic model. All of which are lacking for killer whales and most other non-model species. However, with the development of new genomic tools (e.g. Kim et al. 2017; Huber et al. 2018) and expanding population genomic datasets, the requirements to address such pressing questions in natural population of non-model organisms are becoming increasingly tractable.

## Materials and Methods

Our dataset consisted of twenty-six previously published resequenced 5×coverage genomes (Foote et al. 2019) representative of the known global geographic and genetic diversity of this species. To better understand the temporal context of selection (see Dehasque et al. 2020), we additionally included a 3×coverage genome of an ancient sample from Denmark, radiocarbon dated to 7,530-7,780 years BP (Aaris-Sørensen et al. 2010). Details of generation of the genomic data from the ancient sample and data analyses are given in the supplementary material. Estimates of *L*_*X*/*Y*_ based statistics were estimated as per Do et al. (2015), computing a standard error by jackknifing across the 81 (>10Mb) scaffolds to correct for correlation among neighbouring sites, and using Z-scores to assess significance. The ancestral state for each site was inferred by mapping short reads from a bottlenose dolphin *Tursiops truncatus* (SRX200685) and long-finned pilot whale *Globicephala melas* (SRX5649027) against the killer whale reference genome to a coverage of 5× for each species. Both outgroup species fall into the sister clade to the killer whale, with an estimated Time to Most Recent Common Ancestor (TMRCA) of approximately 10 million years BP (McGowen et al. 2020). The consensus sequence was called using SAMtools (Li et al. 2009), and ambiguous bases were masked with Ns. Coding mutations from autosomal scaffolds >10Mb were annotated from the RefSeq predicted exon coordinates (ftp://ftp.ncbi.nlm.nih.gov/genomes/all/GCA_000331955.2_Oorc_1.1) as ‘non-synonymous’, ‘synonymous’, ‘stop gain’ or ‘stop loss’ and identified as ancestral or derived using ANNOVAR (Wang et al. 2010).

### Ancient DNA extraction, library building, sequencing and mapping

Powdered bone was sampled in a sterile environment from a subfossil killer whale skull fragment of a sample (ZMUC 29/1983) from Lystrup, Jutland, Denmark, which is part of the collections of the Natural History Museum of Denmark (Aaris-Sørensen et al. 2010). The collagen content was estimated at 11.5% from comparison of the weight of the original sample with the weight of the dried collagen prior to radiocarbon dating (Aaris-Sørensen et al. 2010). The sample was radiocarbon dated to 6800±115 years using a conventional proportional counter (Copenhagen Radiocarbon Laboratory ID: K-6800) and calibrated with the OXCAL program using the Reimer et al. (2004) curves to 7,530-7,780 years BP (see Aaris-Sørensen et al. 2010).

Ancient DNA lab work was conducted at the dedicated ancient DNA facilities at the Globe Institute, University of Copenhagen. DNA was extracted from ca. 50 mg bone powder, which was incubated overnight under motion at 55°C in 15 μl extraction buffer (0.45 M EDTA, 0.1 M UREA, 150 μg proteinase K). The sample was then centrifuged at 2300 rpm for 5 min and the supernatant was collected and concentrated and purified using a Zymo-Spin V reservoir (Zymo Research Irvine, CA, USA) and Qiagen MinElute spin column (Qiagen, Inc., Valencia, CA, USA). To maximize library complexity by reducing the number of DNA purification steps during library preparation, an Illumina library was constructed using the blunt-end single tube (B.E.S.T.) method (Carøe et al. 2017). The library was dual indexed and amplified in a 15-cycle PCR using AmpliTaq Gold (ThermoFisher Scientific). The 50μ l PCR reaction contained 15ul of library, 25μ M dNTP, 1x PCR buffer, 2.5 mM MgCl_2_, and was made up to 50μ l with molecular grade water. PCR temperature profile included an activation step at 95°C for 5 min, followed by 15 cycles of denaturation at 95°C for 30 s, annealing at 55°C for 30 s and elongation at 72°C for 1 min, with a final extension step at 72°C for 7 min. PCR products were then purified using the MinElute PCR purification kit (Qiagen). The dual index amplified library had mean insert size of 184bp, including 114bp of index and adapters, indicating mean DNA fragment size of 70bp. Peak molarity was 40,700 pmol/l, comprising 96% of the library (including lower and upper size marker peaks) as quantified using an Agilent 2200 TapeStation instrument with D1000 High Sensitivity ScreenTape and reagents. Extraction, library build and index PCR blanks were also included to evaluate potential contamination during the library building process. DNA libraries were then sequenced across two lanes of 80bp-SE sequencing of an Illumina HiSeq4000 at the Danish National High-throughput Sequencing Centre (seqcenter.ku.dk).

Read trimming, mapping, filtering and repeat-masking was conducted as per Foote et al. (2019). Endogenous content of 10.4% was inferred from the percentage of sequencing reads mapping to the reference genome. The mean length of mapped reads was 57 bp. Of these 62.4% were of base quality ≥30, and of these 79.6% were inferred to be unique reads and were thus used for downstream analyses. In total, 949,721,463 sites were covered at ≥1× coverage (mean per-site coverage of 1.8×) with 86.2% of q-score ≥ 40. Analyses of potential misincorporations using mapDamage2 (Jónsson, et al. 2013) to compare with the modern reference genome revealed that sequencing reads exhibited characteristic post-mortem damage patterns (Dabney et al. 2013), specifically an excess of C>T transitions at the 5′ termini as expected from deamination, and the complementary G>A transitions at the 3′ termini (Supplementary Figure 1). Therefore, only transversions were considered in all downstream analyses that included the ancient sample.

### Inferring the relationship of the ancient genome to modern genomes

To draw inferences on processes such as selection from comparing ancient and modern samples, we must understand how directly ancestral the ancient sample is to our modern samples. Ancient samples from the same location as modern samples need not be directly ancestral, as events such as population extinctions or replacements may have occurred during the time interval in-between. Ideally the ancient sample should be the common ancestor, or at least have experienced minimal drift from the shared most recent common ancestor with the modern sample, so that it represents the ancestral state.

We first generated a time-calibrated killer whale phylogeny using mitochondrial genomes. Thirteen protein coding gene regions were extracted from the 156 modern and one sub-fossil killer whale mitochondrial haplotypes in order to estimate the coalescence times of different killer whale clades. As in Morin et al. (2015), we used only the third codon positions of the genes to avoid any possible effect of incomplete purifying selection on the coalescence times of the tree (Ho, 2005; Ho et al., 2011). The ‘greedy’ search in PartitionFinder (v1.1.0) (Lanfear et al., 2012) was used to find the best partitioning scheme for each gene, and a time-calibrated phylogenetic tree was built in BEAST2 (Drummond et al., 2012) with two different data partitions for nucleotide substitution models (Table S1). We applied a common strict clock model for all of the genes, following Morin et al. (2015). Two independent models were run with 40,000,000 MCMC steps, 10% pre-burn-in, and a sampling frequency of 4,000. The time to most recent common ancestor (TMRCA) of all of the haplotypes (mean of 361.7 kyr and standard deviation of 79 kyr) previously estimated in Morin et al. (2015), was used to calibrate the root of the tree and a Yule prior was used for the tree branching rate. In addition, we applied a tip calibration with the sub-fossil sample carbon-dated to 7,530–7,780 years before present. The convergence of Markov Chain Monte Carlo (MCMC) chains and the Effective Sample Size (ESS) values relating to the model parameters were checked in Tracer (Rambaut et al., 2014). After verifying convergence, LogCombiner and TreeAnnotator (Drummond et al., 2012) were used to combine and summarize the trees, respectively.

The average clock rate for the killer whale haplotypes was estimated as 0.024 substitutions per site per million years (SD = 0.0128), and the TMRCA of all the haplotypes was estimated as 279 kyr (95% HPDI: 84–461 kyr). These estimates are similar to the previous estimates derived in Morin *et al*. (2015). The ESS values for the different parameters in the individual runs were all >600 (most of them >2000), indicating no sign of autocorrelation between samples and a good convergence of chains (Table S2). The summary consensus tree consisting of *O. orca* haplotypes indicates that the ancient sub-fossil sample shares a most recent common ancestor with the basal node to clade 5 which consists of samples collected from the Northeast Atlantic (Supplementary Figure 2).

We then inferred the relationship among ancient and modern samples using the nuclear genome, adopting methods appropriate for low coverage sequence data and DNA damage patterns. We used pseudo-haploid genotype calls of autosomal regions in our globally distributed modern genomes and the ancient sample. Additional filtering steps included the removal of regions of poor mapping quality (Q < 30), removal of sites with low base quality scores (q < 20), calling only SNPs inferred with a likelihood ratio test (LRT) of P < 0.000001, a minimum allele frequency of 0.05 so that alleles had to be called in a minimum of two individuals, discarding reads that did not map uniquely, adjusting q-scores around indels, adjusting mapping quality to 50 for excessive mismatches, discarding bad reads (flag >=256), and the removal of transitions to avoid bias from C®T and A®G DNA damage patterns. Then, we compared covariance among samples in a Principal Component Analysis (PCA) using ANGSD (Korneliussen et al. 2014). The ancient sample was included in the PC computations and not projected onto PCs of modern samples, which has the advantage of providing a quality control measure. For example, if the ancient sample was impacted by sequencing- or sequence data processing errors, the sample would appear as an outlier in the PCA. Instead, the ancient killer whale genome clusters with Atlantic samples in the PCA plot (Figure 2). The eigenvectors from the covariance matrix were generated with the R function “eigen”, and significance was determined with a Tracy-Widom test (Tracy & Widom 1994) performed in the R-package AssocTest (Wang et al. 2020) to evaluate the statistical significance of each principal component.

We then used a likelihood ratio test developed for estimating drift experienced by two populations after their divergence, with no assumptions of the demographic history of either populations. The method was developed for and applied to testing the relationship of ancient human genomes to modern human populations to better understand the peopling of the Americas (Rasmussen et al. 2014, 2015). The model estimates branch lengths of two populations since divergence, each one represented by a single diploid genome, from the probabilities of derived alleles coalescing either along a branch to one of the two populations, or in a shared ancestral population. If the ancient genome is directly ancestral to the modern genome being tested, then the probability of alleles coalescing along the branch to the ancient sample should be low. The method assumes that neither population has admixed with other populations since divergence, an assumption that is potentially violated in our modern samples, and which could overestimate branch lengths to the modern samples in each pairwise comparison. We implemented this method in ANGSD (Korneliussen et al. 2014), removing all transitions and applying additional filters as per steps taken for the PCA and estimating the 2D-Site Frequency Spectrum (SFS) from genotype likelihoods. We infer minimal branch length of the ancient sample in all pairwise comparisons, which while it would be consistent with the ancient sample pre-dating the formation of present-day structure in the North Atlantic, could be downwardly biased by the low coverage of the ancient sample.

To better understand the relationship of the ancient sample to present-day Northeast Atlantic samples, and to investigate whether modern Icelandic and Norwegian genomes evenly shared derived alleles with the ancient samples, we estimated D-statistics (Figure 2). The test can be used to evaluate if the data are inconsistent with the null hypothesis that the tree (((Norway, Iceland), ancient), Gibraltar) is correct. The definition used here is from Durand, Patterson, Reich, and Slatkin (2011):

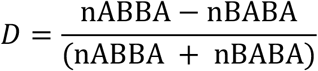

where in the tree given above, nABBA is the number of sites where Iceland and ancient samples share a derived allele, and the Norway sample has the ancestral allele (shared with the Gibraltar sample; and nBABA is the number of sites where the Norway and ancient samples share a derived allele, and the Iceland sample has the ancestral allele. Under the null hypothesis that the given topology is the true topology, we expect an approximately equal proportion of ABBA and BABA sites and thus D = 0. The significance of the deviation from 0 was assessed using Z-scores, which are based on the assumption that the D-statistic (under the null hypothesis) is normally distributed with mean 0 and a standard error achieved using the jackknife procedure. The tests were implemented in ANGSD (Korneliussen et al. 2014) and performed by sampling a single base at each position of the genome to remove bias caused by differences in sequencing depth at any genomic position, removing transitions to avoid bias from C→T and A→G DNA damage patterns, and only considering sites covered in the ancient sample. Further filtering steps were as specified above for PCA. The results are shown in Supplementary Figure 2

## Acknowledgements

A.D.F. was supported by a short visit grant from the European Science Foundation–Research Networking Programme ConGenOmics and by a Swiss National Science Foundation grant (31003A-143393) to L. E., and by the Welsh Government and Higher Education Funding Council for Wales through the Sêr Cymru National Research Network for Low Carbon, Energy and Environment, and from the European Union’s Horizon 2020 research and innovation programme under the Marie Skłodowska-Curie grant agreement No. 663830. Additional costs were covered by an ERC Consolidator Grant 681396 ‘Extinction Genomics’ awarded to M.T.P.G.

## Notes

### Competing Interest Statement

The authors have declared no competing interest.

### Summary of Updates

Figures were wrongly cited in the text.

