## Supplementary Materials for "Evidence of long-term purging of mutation load in killer whale genomes"

**a.**

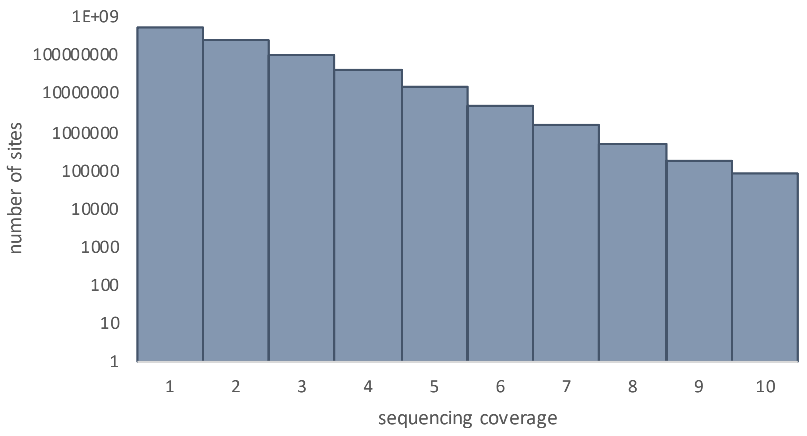


**b.**

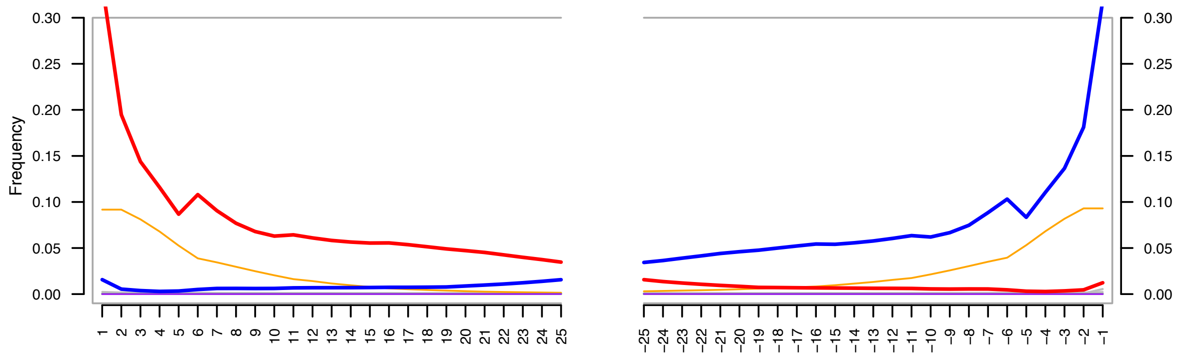


**Supplementary Figure 1.**

**a.** Depth of coverage of the ancient sample, showing up to 10x coverage. **b.** DNA misincorporation errors relative to the 5´ and 3´ read termini sequence data generated from the 7.5KY BP C14-dated ancient sample (29/1983) relative to the modern killer whale reference (Oorc_1.1, GCA_000331955.2). The two distributions for post mortem damage signatures (C>T and G>A) are shown in red and blue respectively. The distribution of soft-clipped bases, *i.e*. bases at the read extremities which did not align with the reference, are shown in orange. These are not taken into account in the computation of damage patterns. The frequency of insertions and deletions relative to the reference sequence are shown in purple and green respectively; all other substitutions are shown in grey. Nucleotide frequencies are shown for 25 bases upstream and downstream of the 5´ and 3´ read termini.

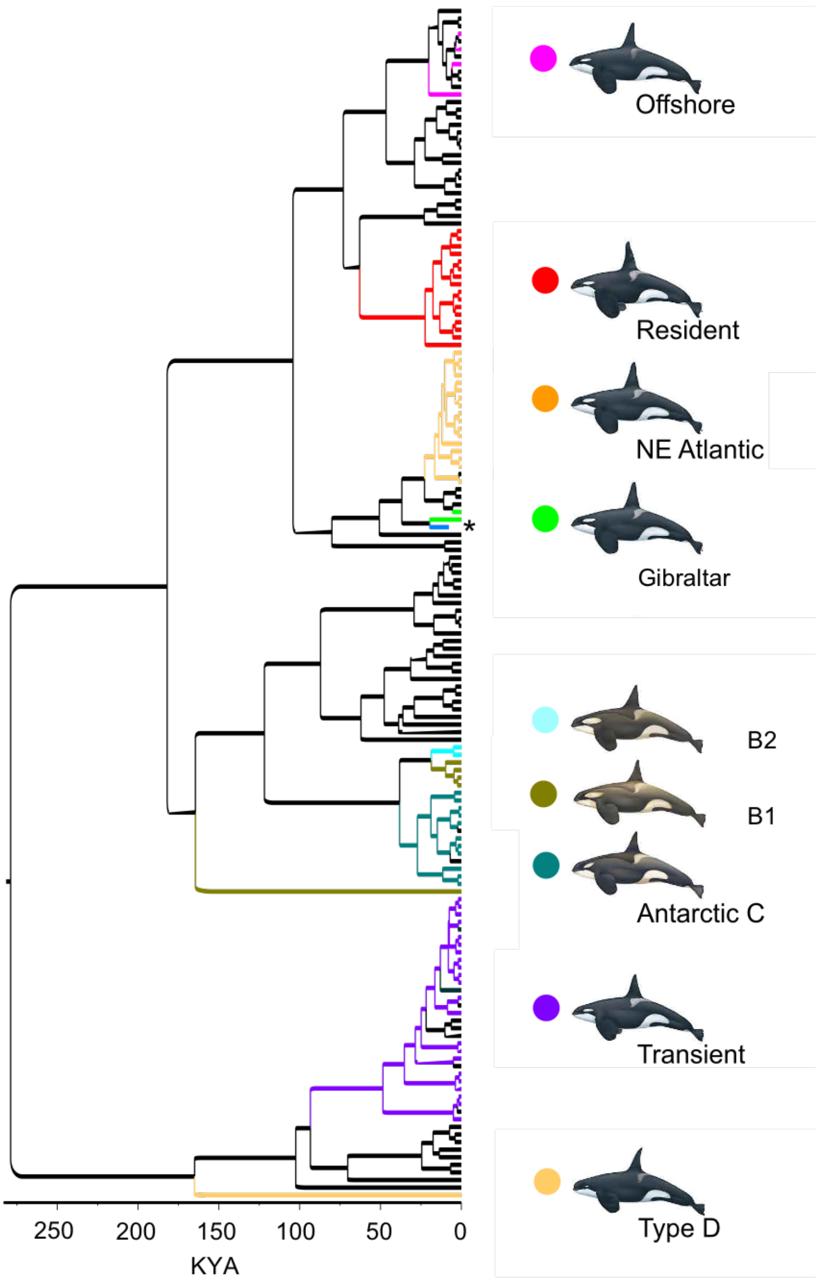


**a.**

**Supplementary Figure 2.**
Bayesian phylogenetic tree of 159 unique haplotype sequences generated from 452 modern samples and one ancient (7,530-7,780 yr BP) sample (marked *), extended from figure 2 in Morin et al. (2015). NE Atlantic clade includes Iceland Norway samples.

Table S1. Best partitioning schemes for nucleotide substitution models used in the construction of the killer whale time-calibrated mtDNA phylogeny

| Partition | Best model | Subset partitions |
|---|---|---|
| p1 | TrN+G | atp6_pos3, atp8_pos3, cox1_pos3, cox2_pos3, cox3_pos3, cytb_pos3, nd1_pos3, nd2_pos3, nd3_pos3, nd4_pos3, nd4l_pos3, nd5_pos3 |
| p2 | HKY | nd6_pos3 |

Table S2. The mean and Effective Sample Size (ESS) values for the different parameters in one run of the BEAST2 time-calibrated models for killer whales. The models were run with a strict clock and including only the third codon positions of the 13 protein coding mtDNA genes.

| Parameter | Mean | ESS |
|---|---:|---:|
| posterior | -6618.95 | 2176 |
| likelihood | -6556.73 | 3564 |
| prior | -62.219 | 2172 |
| treeLikelihood.atp6_pos3 | -407.525 | 855 |
| treeLikelihood.atp8_pos3 | -110.87 | 1692 |
| treeLikelihood.cox1_pos3 | -973.614 | 3475 |
| treeLikelihood.cox2_pos3 | -378.554 | 3965 |
| treeLikelihood.cox3_pos3 | -463.733 | 1013 |
| treeLikelihood.cytb_pos3 | -630.749 | 2478 |
| treeLikelihood.nd1_pos3 | -568.536 | 5394 |
| treeLikelihood.nd2_pos3 | -506.435 | 1948 |
| treeLikelihood.nd3_pos3 | -174.299 | 4641 |
| treeLikelihood.nd4_pos3 | -829.86 | 4580 |
| treeLikelihood.nd4l_pos3 | -170.39 | 3490 |
| treeLikelihood.nd5_pos3 | -1034.41 | 2948 |
| treeLikelihood.nd6_pos3 | -307.758 | 2072 |
| TreeHeight | 0.279 | 2862 |
| clockRate | 0.02144 | 1748 |
| kappa1.1 | 109.176 | 669 |
| kappa2.1 | 50.774 | 671 |
| gammaShape.1 | 0.485 | 2784 |
| kappa.2 | 44.056 | 1578 |
| popSize | 0.541 | 2852 |
| CoalescentConstant | -46.286 | 2156 |
| logP(mrca(Root)) | 0.325 | 2013 |
| mrcatime(Root) | 0.279 | 2862 |
| logP(mrca(Subfossil_tip.prior)) | 8.128 | 1707 |
| height(Subfossil) | 0.007656 | 1804 |
